# Heritable viral symbionts in the family *Iflaviridae* are widespread among aphids

**DOI:** 10.1101/2025.02.03.636351

**Authors:** Paula Rozo-Lopez, Vanesa Torres, Joseph Torres, Simon Käfer, Benjamin J. Parker

## Abstract

Heritable microbes shape host phenotypes and are important drivers of evolution. While interactions between insects and bacterial symbionts have been extensively studied, the prevalence and consequences of insect-viral symbiosis are an open question. We show that viral symbionts in the family *Iflaviridae* are widespread among aphids, an important model for research on bacterial symbiosis. We discovered multiple new species of iflaviruses that are maintained in asexual lines without apparent fitness costs and are transmitted vertically from mothers to offspring. Using field data and phylogenetic evidence, we further show that aphid iflaviruses likely move horizontally across species, but through laboratory experiments, we demonstrated that horizontal transfer among species infesting the same host plants does not persist throughout clonal lineages. Using quantitative PCR and immunohistochemistry, we discovered that viral infections localize in the host fat bodies and developing embryos. Surprisingly, we also found viral infections inside bacteria-housing cells called bacteriocytes, with a positive correlation between viral and bacterial symbiont density, indicating a mechanism for vertical transmission. Together, our work suggests that iflaviruses are an important but previously unrecognized piece of aphid symbiosis and sets the stage to use this model to answer new questions about host-microbe associations.

**Importance:** In recent years, the rise of metatranscriptome sequencing has led to the rapid discovery of novel viral sequences in insects. However, few studies have carefully investigated the dynamics of insect-virus interactions to produce a general understanding of viral symbiosis. Aphids are a significant agricultural pest but also an important model for understanding the evolution of host-microbe interactions and the molecular basis of bacterial symbiosis. We show that heritable iflaviruses are an important but previously unrecognized part of the aphid heritable microbiome, with viral symbionts transmitted alongside bacteria from mothers to offspring, potentially via specialized bacteriocytes that house symbiotic microbes. Our findings have important implications for furthering the understanding of insect-microbe symbiosis and the potential for biocontrol of agriculturally relevant pest species.

## Introduction

The viruses associated with insects have been historically studied within the context of pathogens that directly infect and kill insects and those transmitted to humans, animals, or plants by insect vectors. However, the use of metatranscriptome sequencing over the last two decades has uncovered a vast diversity of novel viruses associated with insects (1–3). Some studies have also shown that specific viral taxa persist across life stages and are inherited through multiple generations (4–6). Though the specificity and persistence of these infections suggest a symbiotic association between viruses and insect hosts (7), little is known about the evolution of heritable viral symbiosis.

In contrast, decades of research have established a clear understanding of interactions between insects and heritable bacterial symbionts. Most insect species interact with heritable bacteria that are passed maternally from reproductive females to offspring (8). Studies stress the energetic costs bacteria impose on their hosts and the need for strategies of vertical spread through host populations either by conferring benefits or with reproductive manipulation (9, 10). Although there is some evidence for host-symbiont specificity, horizontal transmission of bacterial symbionts across insect taxa is a key component of infection dynamics (11, 12). Conversely, these key questions have not been addressed for insect-viral symbiosis. It is unclear, for example, whether heritable viruses coevolve with their hosts, whether they show species specificity with hosts, and how they are maintained in natural populations. To address these questions, the development and study of tractable model insect-virus symbioses are needed.

Aphids (Hemiptera: Aphididae) are arguably one of the most important model insect systems for studying symbiosis (13). Nearly all 5,000 aphid species interact with *Buchnera aphidicola,* an obligate bacterial endosymbiont that provides its hosts with amino acids missing from plant phloem. *Buchnera* and several other species of facultative bacterial symbionts, which are not required for host survival, are vertically transmitted from females to their offspring through asexual (parthenogenetic) and sexual phases of the aphid lifecycle (11, 14, 15). In the aphid, *Buchnera* and some other facultative bacterial symbionts are housed inside specialized cells called bacteriocytes, which play a distinct role in vertical transmission fidelity (16). Recent studies have identified several species of aphid viruses [Bunyavirales (17), *Iflaviridae* (18), and *Parvoviridae* (19, 20)] that seem to be maintained vertically with high fidelity, but little is known about the infection dynamics or the role of aphid-viral symbioses.

The family *Iflaviridae* comprises a single genus (*Iflavirus*) of positive-sense single-stranded RNA (+ssRNA) viruses capable of infecting multiple insect orders such as Lepidoptera, Hymenoptera, Hemiptera, Diptera, Coleoptera, and Orthoptera. In some insects, iflaviruses are potent insect pathogens, while others exhibit asymptomatic persistent infections maintained by a combination of horizontal and vertical transmission (21). Although the genomes of three aphid *iflaviruses* have been sequenced [*Brevicoryne brassicae virus* (BrBV) (18), *Aphis aurantii iflavirus* (Iva) (22), and *Cavariella aegopodii iflavirus 1* (CAIV1) (23)], there is limited information about their biology and transmission dynamics.

Here we found evidence for widespread heritable *Iflavirus* symbiosis across the phylogeny of aphids. We used metatranscriptome sequencing of field-collected aphids and a bioinformatic screen of the NCBI Sequence Read Archive to assemble genomes for multiple new strains and species of heritable iflaviruses. We show that specific viral species can be detected in multiple host species cohabitating on the same host plant, but heritable infections are specific to individual aphid species. Moreover, we demonstrate that the phylogeny of these viral symbionts does not reflect their hosts, suggesting that cross-species horizontal transmission may occur on evolutionary timescales. We also show that *Iflavirus* infections have no clear negative fitness effects on aphids and are present in specific tissues like fat bodies, ovaries (developing embryos), and bacteriocytes, which could explain the high infection rates of these viruses. Lastly, we show that the density of *Iflavirus* infections correlates with bacterial symbiont titers, suggesting an interaction between these two microbes that could be key to future work. We discuss the potential implications of long-term symbiotic interactions between viruses and their hosts while building a comprehensive picture of heritable virus-host dynamics in an important insect group.

## Results

### *Iflavirus* discovery using RNAseq

We described several new species of iflaviruses associated with aphid hosts through metatranscriptome sequencing of field-collected aphids. Initially, we established laboratory colonies from field-collected parental aphids (P) of the tribe Macrosiphini (Wilson, 1910) and maintained them for at least three subsequent generations (F_1_–F_3_) (Table S1). RNAseq data was derived from ribosomal RNA-depleted libraries created for a subset of the parental lines (Table S2) and analyzed using two methods. First, we utilized CZ ID, a cloud-based open-source bioinformatics platform, to screen for viral reads matching iflaviruses and unclassified RNA viruses (Fig. S1). Second, we employed TRAVIS, a consistency-based virus detection pipeline for sensitive mass screening of transcriptomic data focused on *Iflaviridae* proteins, to obtain long viral contigs (>10,000 nt). We selected assemblies that exhibited nucleotide and amino acid similarities to members of the family *Iflaviridae* (9–11 kb genomes encoding a single large polyprotein) and used the aphid *Brevicoryne brassicae virus* (BrBV; GeneBank accession YP_001285409.1) as a reference to generate consensus genomes. Following the ICTV species demarcation criteria for iflaviruses, we classified as new viral species genome assemblies obtained from different host species that showed a capsid protein (VP) amino acid sequence similarity below 90% (24). Within the same virus species, we considered genome assemblies from distinctive geographical locations as new viral strains. This approach produced complete genomes (99.9% coverage breadth) for a new strain of BrBV (genome size of 10,179 nt) and three new *Iflavirus* species, which we named *Ilinoia iflavirus* (IspV; 10,546 nt)*, Uroleucon eupatarifoliae virus* (UeV; 10,483 nt), and *Uroleucon iflavirus* (UspV; 10,340 nt) (Fig. 1a; Table S3).

**FIG 1.**
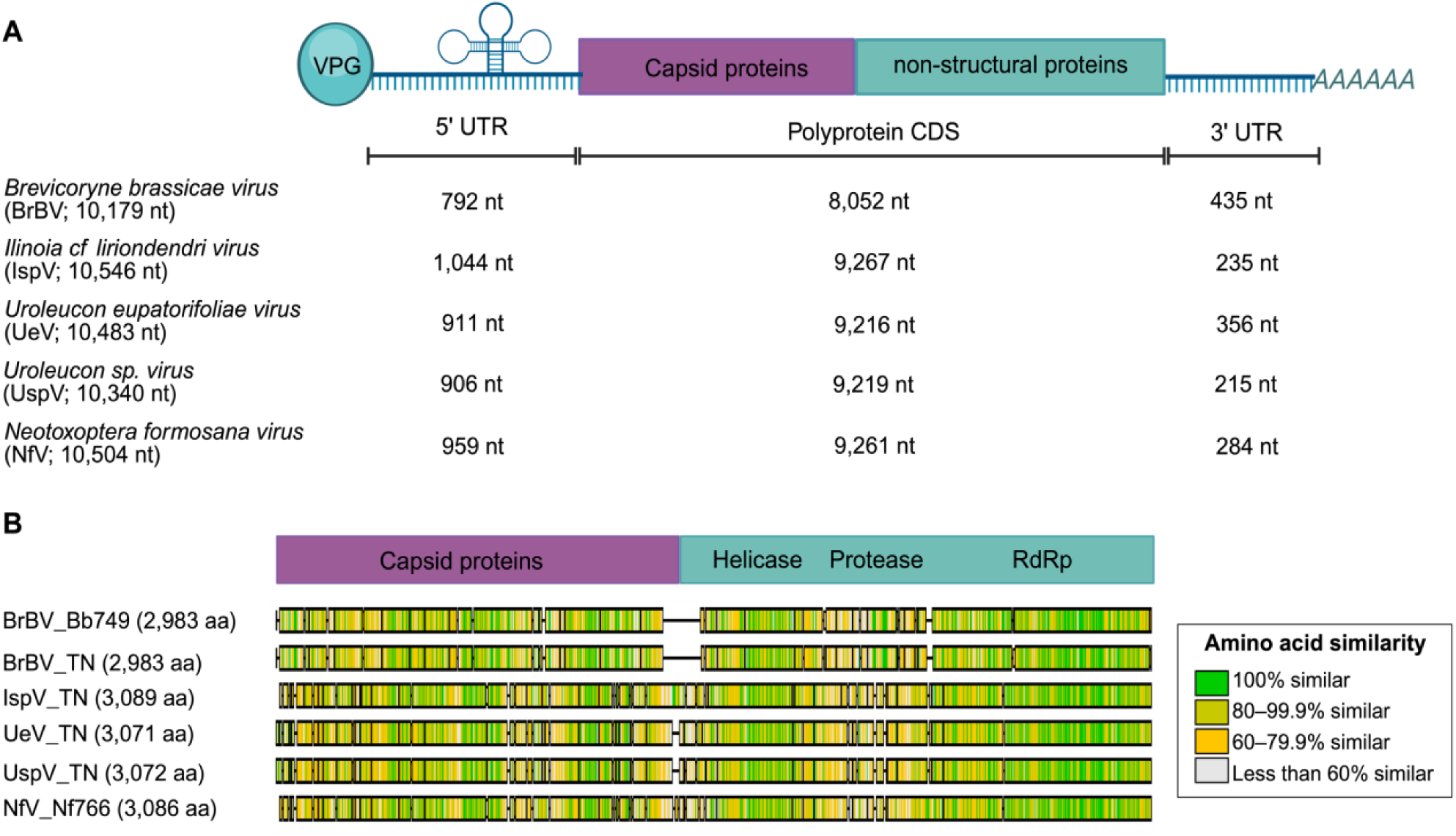
Newly identified aphid iflaviruses. (**A**) Complete consensus genome structure specifying the nucleotide (nt) lengths of the polyprotein coding region and the 5′ and 3′ untranslated regions (UTR). (**B**) Alignment of polyprotein sequences showing the amino acid (aa) similarity in color scale. Green colors denote higher similarity per site, yellow moderate, and grey low similarity. BrBV_TN, IspV_TT, UeV_TN, and UspV_TN correspond to the viral sequences obtained from metagenomic data of field-collected insects. BrBV_Bb749 and Nf_N766 correspond to the viral sequences obtained from SRA datasets.

### *Iflavirus* discovery in the Sequence Read Archive

In addition to our sequencing efforts, we identified further evidence of widespread *Iflavirus* prevalence in existing datasets within the Sequence Read Archive (SRA). After screening 27 RNAseq datasets (22 species) (Table S4), we assembled another sequence of BrBV, UK strain (genome size of 10,179 nt), along with a new *Iflavirus* species that we named *Neotoxoptera formosana virus* (NfV; 10,504 nt) (Table S3). We also obtained a 7,092 nt sequence of the coding region from a potentially new *Iflavirus* hosted by *Megoura crassicauda* (named *Megoura crassicauda virus*; McV), but it was not included in our further analysis due to the absence of a complete polyprotein sequence.

All six newly assembled *Iflavirus* genomes have a single large ORF (CDS 8,952–9,267) flanked by a 5′ untranslated region (5′ UTR), which includes an internal ribosome entry site (IRES, required for the cap-independent translation) and a 3′ UTR with a poly(A) tail that terminates translation (21) (Fig. 1A). The amino acid similarity of the newly assembled aphid Iflavirus polyproteins is the highest at the RNA-dependent RNA polymerase (RdRp) region (Fig. 1B), while the overall polyprotein similarity ranging from 33.32 to 87.96% (Table S5).

### *Iflavirus* prevalence in field samples

We found that individual *Iflavirus* species can be detected via PCR in multiple hosts and are found at intermediate frequencies within species. We tested the *Iflavirus* prevalence across the field-collected parental (P) aphid lines using multiple species-specific primer sets (Table 1). We detected BrBV in both field-collected *B. brassicae* (72.2%) and *Lipaphis pseudobrassicae* (16.7%) infesting *Brassica* plants. Similarly, we found UeV with 100% prevalence in *Uroleucon eupatoricolens* and in a percentage (20%) of different morphotypes of aphids from the genus *Uroleucon* (large daisy aphids) whose species assignment could not be confirmed due to the lack of progeny to obtain functional barcode sequences. Likewise, UspV infections ranged from 0–100% across multiple species of aphids in the genus *Uroleucon* infesting Asteraceae. We also detected iflaviruses that infected only a single aphid host species, as in the case of IspV found in 22.2% of *Ilinoia liriodendri* aphids infesting tulip trees.

**TABLE 1.** *Iflavirus* prevalence and heritability in aphid colonies.

| *Iflavirus* | Aphid species | Percentage of *Iflavirus*-positive colonies |  |
| --- | --- | --- | --- |
|  |  | Parental (P) | Progeny (F<sub>1</sub>–F<sub>3</sub>) |
| BrBV | *Brevicoryne brassicae* (Bb) | 72.2% (13 out 18) | 100% (11 out 11) |
|  | *Lipaphis pseudobrassicae* (Lp) | 16.7% (1 out 6) | 0% (0 out 4) |
| IspV | *Ilinoia liriodendri* (Il) | 22.2% (4 out 18) | 100% (3 out 3) |
| UeV | *Uroleucon eupatoricolens* | 100% (3 out 3) | 100% (2 out 2) |
|  | *Uroleucon* spp. | 20% (1 out 5) | Nc* |
| UspV | *Uroleucon eupatoricolens* | 33.3% (1 out 3) | Nc* |
|  | *Uroleucon nigrotuberculatum* | 0% (0 out 1) | Nc* |
|  | *Uroleucon rurale* | 100% (2 out 2) | 100% (2 out 2) |
|  | *Uroleucon* spp. | 100% (5 out 5) | Nc* |

\*Nc: not colonized; aphids died before reaching three generations in the laboratory

### *Iflavirus* maintenance in laboratory colonies

We showed that *Iflavirus* infections were maintained in laboratory lines and vertically transmitted from females to offspring across at least three generations; a critical benchmark for determining heritability (25). We assessed *Iflavirus* heritability in laboratory colonies of the virus-positive parental strains by PCR testing individual adults (1–3) after one (F_1_) and three (F_3_) generations after laboratory colonization (Table 1). All four *Iflavirus* species were maintained with 100% fidelity in the laboratory. BrBV and IspV further showed stable heritable infections for more than 30 generations (over one year) and, therefore, were used for subsequent transmission experiments.

### *Iflavirus* vertical transmission in lab colonies

We found evidence that vertical transmission of BrBV and IspV occurs from mothers to offspring. All *B. brassicae* (Bb) and *I. liriodendri* (Il) females vertically transmit their corresponding iflaviruses to their offspring (vertical transmission rates; VTR= 100%), but the proportion of infected offspring (collected immediately as they were being born from a single *Iflavirus*-positive female) varied across species and lines (filial infection rates; FIR). In Il aphids, the FIR was 84.6% (±0.11), and in Bb aphids, the FIR ranged from 98.7% (±0.2) to 43.7% (±0.11), depending on the line. We did not find significant differences (Mann-Whitney test, p = 0.9591) in relative BrBV adult body titers (qPCR) that could explain the difference in FIR between lines (Fig. 2A). In an attempt to produce a parthenogenic BrBV(−) line, we split the line with the lowest FIR into multiple sublines and maintained them for five generations at lower aphid densities while randomly testing 1–3 adults at every generation. Although we observed some PCR-negative adults, our qPCR analysis revealed that those lines still retained the virus at titers below the PCR limit of detection, and BrBV was detectable again by PCR when aphids were subjected to crowded conditions. Together, these data suggest that aphid iflaviruses are highly heritable.

**FIG 2.**
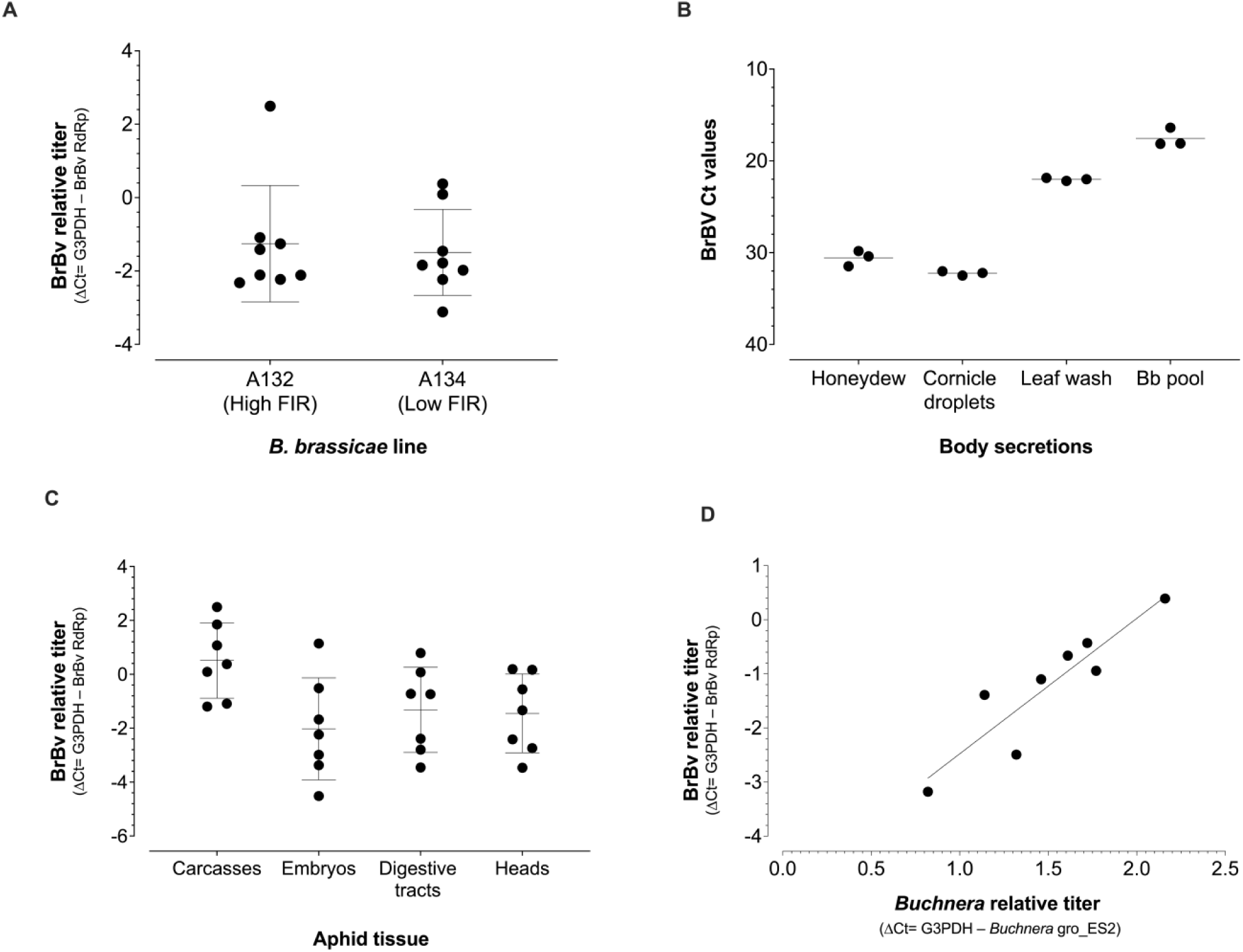
*Iflavirus* infection titers (qPCR). (**A**) Relative BrBV titers in *B. brassicae* adults sampled from the lines with the lowest and highest filial infection rates (FIR). (**B**) BrBV Ct values in honeydew, cornicle droplets, and PBS washes of plant surfaces contaminated with body secretions. Pooled adult *B. brassicae* adults (BB pool) from which honeydew and cornicle droplets are used as a reference positive. (**C**) Relative BrBV titers in organs dissected from *B. brassicae* adults. (**D**) Correlation between BrBV and *Buchnera* relative titers. (A, C, D) The aphid reference gene G3PDH was used to normalize the data.

### Within species horizontal transmission of iflaviruses

We found that intraspecific horizontal transmission contributes to the maintenance of aphid iflaviruses through cohabitation but not via viral ingestion. We observed that 100% of F_1_ adults were PCR-positive for BrBV and IspV infections when allowed to cohabitate with *Iflavirus*-positive reproductive females for four days on leaf cuttings before being transferred to new host plants. We also found that honeydew and cornicle droplets collected from Bb adult aphids and PBS washes of plant surfaces contaminated with body secretions were qPCR-positive for BrBV (Fig. 2B), indicating that transmission of BrBV during cohabitation is enhanced by virus-positive body secretions.

Regarding oral infections, we found that 100% of Il aphids (from the *Iflavirus*-negative line) were PCR-positive for IspV (n=5 pools) right after feeding for 24 hours on artificial diets supplemented with IspV(+) aphid homogenates. However, none of the aphids tested positive for IspV infections after two or seven days after being placed in virus-free plants (n=5 and 3 pools, respectively), suggesting that ingested virions do not produce active viral infections. Likewise, 4.8% of nymphs were PCR-positive for IspV (1 out of 21) after feeding for 3 days on IspV homogenates delivered through the host plant roots. As before, none of the IspV-fed aphids that reached adulthood on virus-free plants (n=18) or their progeny (n=18 pools) were positive for IspV infections, indicating that *Iflavirus* spread through the plant phloem does not result in host heritable infections.

### Cross-species transmission of iflaviruses

Through a series of experiments, we found that aphid iflaviruses are species-specific and do not produce heritable infections in distantly related aphids cohabitating on *Brassica* plants. Driven by our field data, we explored if BrBV is horizontally transmitted between *L. pseudobrassicae* [Lp; BrBV(−)] and *B. brassicae* [Bb; BrBV(+)] infesting the same host plants. Initially, we found that 75% (9 out of 12) of Lp aphids were PCR-positive after cohabitation with BrBV(+) aphids on leaf cuttings. After transferring those BrBV(+) Lp adults to virus-free plants in the absence of virus-positive aphids, we observed that all the Lp offspring produced were negative for BrBV infections (qPCR; n=31), suggesting that the Lp reproductive females tested positive for BrBV after cohabitation due to surface contamination with body secretions rather than active heritable infections.

When we tested the potential of BrBV oral cross-species transmission through the host plant, we also found that 50% (2 out 4) of Lp aphids were PCR-positive for BrBV after the initial feeding on the virus-plant interface (before the gut-clearing step, which removes ingested virus that is not acquired) but did not find any virus-positive Lp aphids after reaching adulthood on virus-free plants (qPCR; n=29). We were unable to use artificial diets supplemented with homogenates from the BrBV(+) lines in Lp aphids since we could not verify the feeding status due to their darker green cuticle color. Overall, our data suggest that BrBV oral infections do not result in Lp active infections.

Lastly, when BrBV was microinjected to bypass all infection barriers, we detected that 100% of aphids were PCR-positive for BrBV right after injection, but only 48% were positive three days post injection (3 dpi). We did not detect BrBV(+) results in microinjected Lp after 7 dpi or in pools of nymphs produced by injected reproductive females, which indicates that BrBV has a highly specific host range and the *L. pseudobrassicae* parental aphids testing positive for BrBV in our field survey represent cross-contamination of viral RNA via body secretions rather than active infections.

### *Iflavirus* tissue tropism

We found that BrBV is widely distributed across aphid tissues and with infection patterns relevant to vertical transmission. We detected BrBV (qPCR) in dissected *B. brassicae* heads, digestive tracts (with bacteriocytes attached), developing embryos, and carcasses (with fat body attached), with carcasses showing the highest relative viral titers, although not statistically significant (Kruskal-Wallis test, p = 0.0729; Fig. 2C). It is important to note that material for the tissue examination was pooled from several individuals and could be possibly cross contaminated to some degree by body secretions released during the dissection process. For a more precise description of *Iflavirus* tissue tropism, we used immunohistochemistry to visualize the BrBV infection patterns on longitudinal sections of adult *B. brassicae* (Fig. 3). We observed positive intracellular staining in 81.8% of the examined aphids (18/22), with fat bodies being the most prevalent positive tissue (positive signal detected in 72.7% of aphids examined) (Fig. 3C, E, K), followed by developing embryos (63.6%) (Fig. 3C, E, I), and bacteriocytes (22.7%) (Fig. 3C and G). No positive staining was observed in the brain, hindgut, or midgut suggesting these tissues may be refractory to BrBV infection.

**FIG 3.**
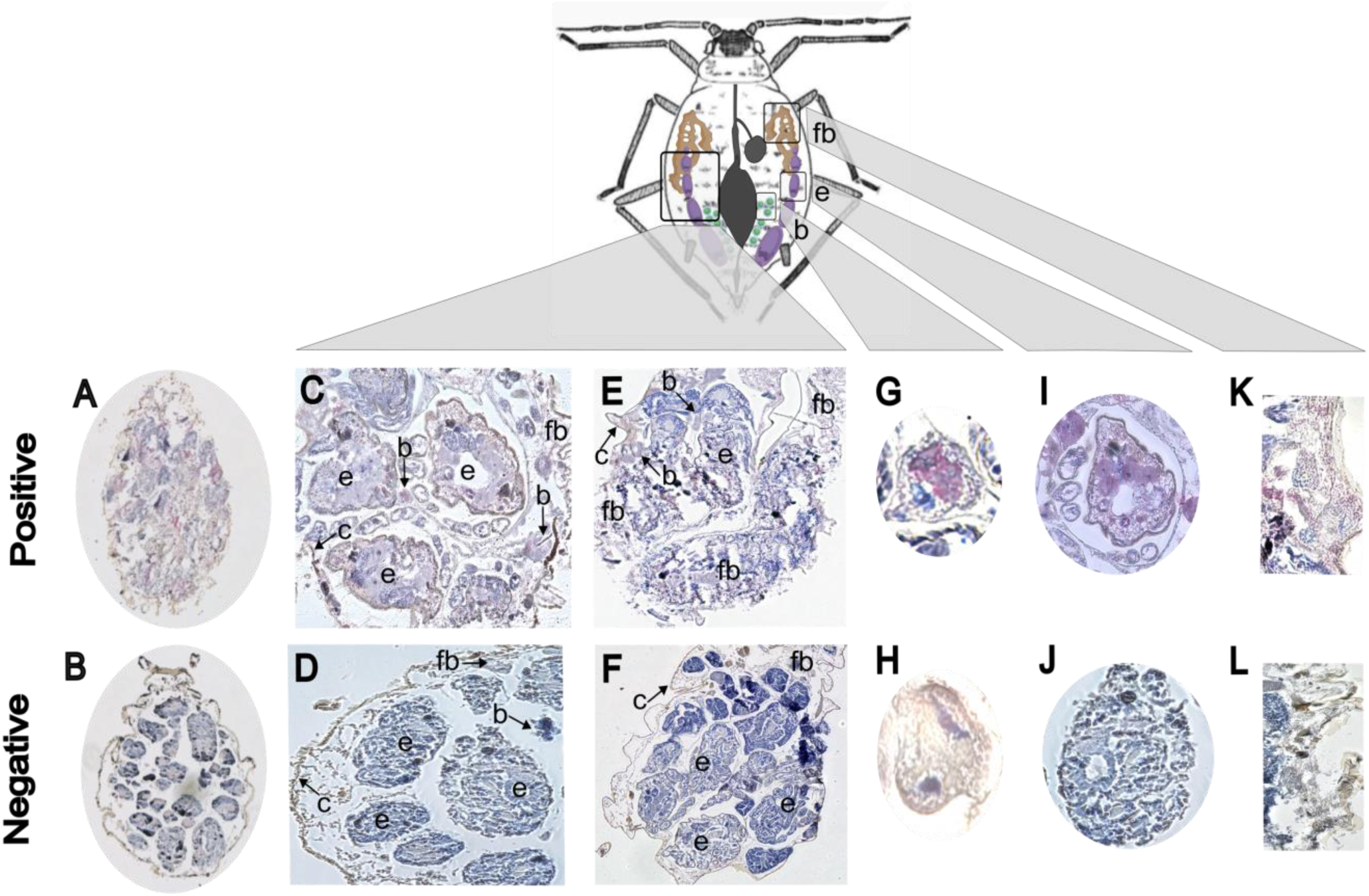
Immunohistochemistry staining of BrBV infected aphid tissues. (**A**) Hematoxylin-stained longitudinal sections of *B. brassicae* and (**B**) *L. pseudobrassicae*. (A, C, D, E, G, I, K) Longitudinal sections of *B. brassicae* tissues showing BrBV-positive (pink) staining. (B, D, F, H, J, L) Longitudinal sections of *L. pseudobrassicae* tissues were used as negative-staining controls. (**G**) BrBV-positive and (**H**) BrBV-negative bacteriocyte. (**I**) BrBV-positive and (**J**) BrBV-negative embryo. (**K**) BrBV-positive and (**L**) BrBV-negative fat body. All sections were counterstained with hematoxylin (blue) to reveal the overall tissue structure. Abbreviations: b: bacteriocyte, c: cuticle, e: developing embryo, fb: fat body.

### *Iflavirus* influence on aphid fitness

We observe that iflaviruses are maintained in laboratory lines without direct fitness cost to aphids. We used IspV(+) and IspV(−) *I. liriodenri* lines to evaluate the effects of iflaviral infection on aphid’s life history traits, although it is important to note that those are not genetically identical aphid lines. We did not find any significant changes in fecundity (Mann-Whitney test, p = 0.4502), with a single IspV(+) parental female producing on average 9.33 (± 4.13) babies and an IspV(−) female producing 7.5 (± 2.42) offspring. Likewise, neither the developmental time (12.5 ± 0.84 days for both lines; p > 0.999) nor stress-induced wing production after starvation and crowding (0% of winged offspring for both lines; p > 0.9999) changed with virus status (Fig. S2). Our results suggest that symbiotic *Iflavirus* infections seem not to negatively influence the fitness of aphids maintained in the laboratory. In addition, we measured the within-host density of the obligate bacterial symbiont *Buchnera* across *B. brassicae* aphids with variable BrBV titers and found a positive correlation between relative titers of bacterial and viral symbionts (Fig. 2D; Spearman Rank-Order correlation, r = 0.9048, p = 0.0023). One possibility is that this trend is driven by the colocalization of both microbes in bacteriocytes, with bacteriocyte numbers potentially differing across generations (26).

### Phylogenetic analysis of aphid iflaviruses

Despite the apparent species specificity of aphid iflaviruses, we found evidence of horizontal transmission across the aphid phylogeny on evolutionary timescales. In a Maximum-likelihood tree based on a MAFFT alignment of the polyprotein sequences of all available aphid iflaviruses (Table S6) and 37 reference species, we found that all aphid iflaviruses cluster together and have other Hemipteran iflaviruses as closely related groups (Fig. 4A). Within the aphid Iflavirus clade, there are two distinctive groups with 100% bootstrap support. Group 1 contains all BrBV strains, CAIV1, and IaV; whereas group 2 includes *Uroleucon* iflaviruses (UeV and UspV), IspV, and NfV. We are confident of the monophyletic nature of aphid Iflaviruses, as well as their clustering patterns, since they were reproducible by other methods such as neighbor-joining (Fig. S2) and even when we included an additional 221 curated sequences from a wide range of iflaviruses infecting arthropods (Fig. S3). Likewise, across all phylogenies, we observed a clade of planthopper honeydew viruses (*Nilaparvata lugens* and *Laodelphax striatellus* honeydew viruses) as the closest related group to the aphid *Iflavirus* clade. Lastly, when comparing the aphid *Iflavirus* phylogeny with the most recent molecular phylogeny of Macrosinphini (27), we found that the virus phylogeny does not fully recapitulate the same clustering patterns of aphid hosts (Fig. 4B), potentially due to disruptive events such as host switching and/or horizontal transmission across host species.

**FIG 4.**
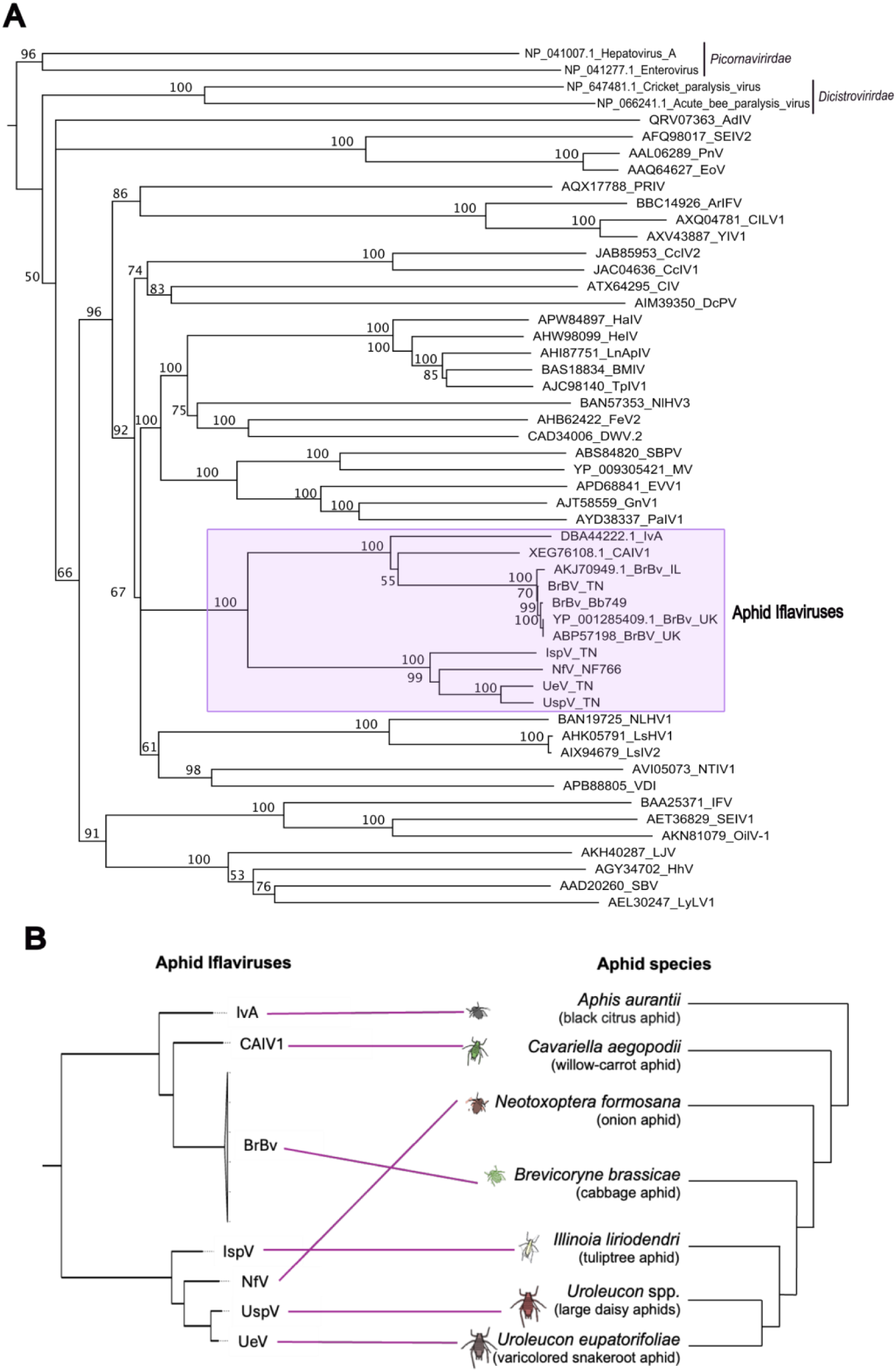
Evolutionary analysis of aphid iflaviruses. (**A**) phylogenetic tree for aphid iflaviruses and ICTV-recognized members of *Iflaviridae*. The tree was constructed using the Maximum-likelihood method with the protein gamma JTT matrix model based on an MAFFT alignment (FFT-NS-i_x1000, BLOSUM62, and 1.54 gap penalty). Families *Dicistroviridae* and *Picornaviridae* are used as outgroups. Bootstrap values on each node represent the percent from 500 replicates. (**B**) representation of phylogenetic trees of aphid iflaviruses (left) and aphid species (right; modified from (27)). Purple lines indicate which hosts harbor a particular *Iflavirus*.

## Discussion

In this study, we integrated metatranscriptomic and molecular screening to investigate the prevalence of novel iflaviruses in various aphid species, with a thorough examination of the dynamics of these heritable infections. Our results revealed that iflaviruses have a long history of interaction with aphids and are a crucial component of these insects’ heritable microbiome, where maternal and horizontal transmission serve as key drivers of viral spread throughout aphid populations and across evolutionary timescales. Our work paves the way for using this established model of bacterial symbiosis to explore new questions about the evolution of insect–virus symbioses.

Insect iflaviruses have been widely studied due to their pathological effects on keystone species such as honeybees (28–30). However, in most insects, infections are asymptomatic and usually detected through next-generation sequencing (31, 32), as is the case of the three species of iflaviruses previously described in aphids (18, 22, 23). Here, we described four additional novel *Iflavirus* species present in several host species (Fig. 1, Table 1), indicating that heritable symbiosis with Iflaviruses is widespread across the aphid phylogeny. In addition, our laboratory experiments showed these heritable infections are maintained as persistent species-specific infections with fluctuating titers (Fig. 2). Although cross-species replication of symbiotic *iflaviruses* has been reported in other insects (31, 33, 34), our cross-species experiments indicate that at least BrBV does not produce active infections in distantly related aphids (*L. pseudobrassicae*) cohabitating on *Brassica* plants with virus-positive hosts (*B. brassicae*). Still, our field survey indicated that UspV could be infecting several species of *Uroleucon* aphids (Table 1), but further analysis is needed to determine if *Uroleucon* iflaviruses are capable of replication in multiple closely related hosts or if there could be different viral strains infecting specific hosts as a sign of an early process of diversification.

Regarding transmission, many viral symbionts, including several iflaviruses, combine vertical and horizontal spread (21, 35), but the mechanisms underlying the horizontal transfer vary greatly among microbe–host pairings. For example, the oral route seems to be the most efficient transmission mechanism for the brown leafhopper *Iflavirus Nilaparvata lugens honeydew virus-1* (36), whereas transmission by a vector (*Varroa destructor* mite) is required for *Deformed wing virus* transmission between bee colonies (37–39). In the case of aphid iflaviruses, we found that horizontal transmission occurs mainly due to contamination with body secretions (honeydew and cornicle droplets), which are commonly encountered during co-habitation. In other Hemipterans, it has been shown that infections with body secretions occur when viral particles of *Euscelidius variegatus iflavirus 1* penetrate the cuticle of leaf hopper nymphs, pass through the tracheae, and then replicate in the insect body (40).

Other aphid heritable viruses (*Parvoviridae*: *Densovirus*) have been shown to systemically spread through the plant phloem after saliva injection (20, 41). Although BrBV replication does not occur in the plant leaves hosting infected aphids (18), we detected relatively high *Iflavirus* loads on plant surfaces (Fig. 2C). Therefore, we conducted several feeding assays to explore if ingestion of iflaviruses could result in systemic heritable infections. We found that iflaviruses can travel through plant tissues, and oral ingestion occurs within 1-3 days, but virus-positive aphids typically exhibit low viral titers that quickly fade after a gut-clearing period. Since ingested virions do not produce active heritable infections and we did not observe positive *Iflavirus*-staining in the digestive tract, we presumed that there might be a midgut infection barrier in which *Iflavirus* particles cannot infect the midgut epithelial cells and/or disseminate to the hemocoel.

The high heritability rates observed in our experimental assays (Table 1) could be favored by laboratory-rearing conditions (e.g. shared housing environments) that may increase virus titers and transmission opportunities through postoviposition secretions (18, 42, 43). However, our careful examination of tissue infection patterns inside developing embryos and bacteriocytes (Fig. 3) confirmed that aphid iflaviruses are vertically transmitted from mothers to offspring. Although *Iflavirus* tissue infection varies among host-virus pairings, ovarian tissue infection seems to be a feature common to iflaviruses infecting Hemiptera (40, 44). While we still need to conduct further investigations to determine the exact transfer mechanisms by which iflaviruses are incorporated into the developing embryos, the positive viral staining inside the bacteriocytes suggests that aphid iflaviruses could co-opt essential maternal transmission mechanisms used by obligate and some facultative bacterial symbionts (45). In aphids, *Buchnera aphidicola* cells are delivered from bacteriocytes (which cluster together near the germ cells) directly to the oocyte via cell-to-cell transfer from the follicle cells during sexual reproduction and via exo/endo-cytosis during parthenogenetic embryogenesis (46). Alternatively, virus particles can bind their capsids to the outer membrane of the obligate symbionts to exploit their oocyte entry pathway (47).

We also localized iflaviruses in the fat bodies (shaped in two bands extending down the lateral abdomen), an important endocrine tissue essential for biosynthetic and metabolic activity that offers a suitable niche for the proliferation of endosymbionts (48). It has been shown that multiple bacterial and viral endosymbionts reside in the insect host fat body (32, 49–52), with some infections altering their host glucose and glycogen metabolism via insulin signaling (53). Additionally, the fat body plays a crucial immune function during viral infection (54), and the high load of BrBV (positive staining and carcass qPCR titers) detected in the fat body might be contributing to the immune homeostasis necessary for maintaining persistent infections (55, 56).

We did not find any apparent detrimental effects of symbiont infection on the aphids’ fecundity, developmental times, or stress-induced wing production, but future studies are needed to evaluate potential context-dependent effects in the parasitism-mutualism continuum. Moreover, in viruses exhibiting wider host ranges, as in the case of potential cross-species transmission of *Uroleucon* viruses, increased virulence due to maladaptation in the newest host is possible, which might suggest that the asymptomatic infections only occur in the original host species (57).

Lastly, our phylogenetic analysis showed a punctuated distribution of aphid iflaviruses, indicating that this host-virus association could have been ancestral to the taxonomic group. While vertical transmission of symbionts leads to a shared reproductive fate that reflects the symbiont phylogeny broadly mirroring the host phylogeny, horizontal transfer events can lead to incongruent relationships (58). Our phylogenetic results (Fig. 4) showed a phylogenetic congruency in *Uroleucon* and *Ilinoia* aphids and their viruses, but the phylogeny of *Brevicoryne* and *Neotoxoptera* iflaviruses did not recapitulate the phylogeny of their aphid hosts. Given that horizontal virus transmission across species is the dominant mechanism in the evolution of RNA viruses (59), random events of horizontal transfer between hosts sharing similar habitats (as suggested by our field screening results) could result in evolutionary events leading to closely related *Iflavirus* species occurring in not-so-closely related hosts. Also, resolving discrepancies in the aphid phylogeny (60) and increasing the virus-host sampling size could improve the analysis resolution and reveal additional evolutionary patterns in the aphid-*Iflavirus* relationships.

Overall, our study of heritable *Iflavirus* infections in aphids indicates that viruses are key players in insect-symbiont interactions and co-evolution. Future studies on the epidemiology and phylogeny of other symbiotic viruses in nature, combined with laboratory experiments, will help improve the general understanding of viral symbiosis. Untangling the factors that influence host-microbe interactions is also crucial for improving knowledge of host-microbe symbiosis in emerging systems and might even have further implications for the prediction and mitigation of emerging viral infections and the control of agricultural pests.

## Materials and Methods

### Aphid collection and colonization

We collected asexual winged and wingless female adult aphids infesting different host plant species across urban and agricultural landscapes in East Tennessee, USA, between April–June and October–December 2022–2024 (Table S1). We placed single adult aphids from the field in Petri dishes containing a leaf disc of the appropriate host plant and allowed them to reach parthenogenic reproduction in the laboratory. Then, individually collected surviving parental adults (P) that had produced offspring (n>5) and stored each parental aphid individually in Eppendorf tubes at −80°C until further processing. We transferred generation one (F_1_) nymphs (offspring produced in Petri dishes) onto laboratory-grown, virus-free host plants and reared them under standard conditions (22°C, 16L:8D, and 45% relative humidity). Every two weeks, we transferred a new generation of nymphs onto fresh laboratory-grown plants (see Table S7 for host plant details), collected 1–3 adults per generation in Eppendorf tubes, and stored them at −80°C until further processing. We identified each aphid species as previously reported (61, 62) by DNA extraction, PCR amplification, and Sanger sequencing of the COI barcode region. Barcoded sequences are available in NCBI with accession numbers PV016919– PV01693. Aphid line metadata is available in Table S7.

### Total RNA extraction

For all RNA extractions, we used −80°C-frozen aphids, tissues, body secretions, or homogenates for pestle-maceration in 300 µL of TRIzol (Invitrogen; Thermo Fisher Scientific, Inc., Waltham, MA, USA) with 100 μL BCP (1-bromo-3-chloropropane; Life Technologies, Thermo Fisher Scientific, Inc., Waltham, MA, USA) and standard isopropanol precipitation. We resuspended each RNA pellet in 40 μL nuclease-free water, removed the genomic DNA with Zymo DNase I Reaction Mix according to the manufacturer instructions (Zymo Genetics Inc., Seattle, WA, USA), and cleaned and concentrated with the Zymo RNA Clean & Concentrator kit (Zymo Genetics Inc.) under recommended conditions.

### Metatranscriptome sequencing

We pooled equal amounts of RNA from between 1 and 6 aphids (P) of the same species (Table S2) to perform metatranscriptome sequencing at Novogene (Novogene Corporation Inc., Sacramento, CA, USA). Library preparation was conducted using ribosomal RNA (rRNA) depletion by Illumina TruSeq Stranded Total RNA with Ribo-Zero Plus and NEBNext rRNA Depletion Kit. Libraries were sequenced to approximately 9 billion base pairs (bp) per sample with 150 bp paired-end reads on an Illumina NovaSeq platform. Raw reads were deposited into the NCBI Sequence Read Archive under BioProject ID PRJNA1216975, accession numbers SRX27542651– SRX27542654.

### *Iflavirus* detection in RNAseq data

We used the CZ ID platform (https://czid.org) with Illumina mNGS Pipeline v8.2 (63, 64), removed aphid-specific reads using the *Acyrthosiphon pisum* genome as host (65), and added The CZ ID water background model after processing. We identified the potential viral reads using z-score metrics (≥1), alignment length over 50 matching nucleotides (NT L ≥50), and a minimum of five reads per million aligning to the reference protein database (NR rPM ≥ 5) (63, 66, 67). Viral contigs were confirmed with manual searches using BLASTX and BLASTN (NCBI). In addition, non-assembled reads matching *Iflaviridae* or unclassified RNA viruses (as a proxy to capture putative Picorna-like viruses with potential *Iflavirus* polyprotein motifs) were manually downloaded and used to perform a *de novo* assembly using Geneious Prime v.2023.2.1 (https://www.geneious.com) at medium sensitivity. We conducted an additional screening for potential *Iflavirus* open reading frames (ORFs) using TRAVIS (v.20221029, https://github.com/kaefers/travis) on *de novo* transcriptome assemblies generated on Trinity v.2.15.1 using default settings (68). Our *Iflaviridae* reference database included the accepted viral species by the International Committee on Taxonomy of Viruses (ICTV) by 24 November 2023 (Table S8). Potential *Iflavirus* ORFs (100–3,000 amino acids) were extracted from the assembled transcriptomes and screened using HMMER v3.3.1 (69), MMSeqs2 (70), BLASTP v2.12.0 (71), and Diamond v2.0.15 (72) with an e-value cutoff of 1 ×10^−6^. All hits were again searched with Diamond against the NCBI non-redundant protein database.

### *Iflavirus* genome analysis, coding region, and polyprotein sequence

For *Brevicoryne brassicae virus* (BrBV), we used the CZ ID viral consensus genome pipeline to build a consensus genome by aligning the *Iflavirus* reads to the reference BrBV genome (NCBI accession NC_009530.1). For the other *Iflavirus* genomes, we aligned CZ ID and *de novo* assembled *Iflavirus* and Picorna-like contigs (>1,000 nt) with the viral contigs provided by TRAVIS using the Geneious Prime Culstal Omega v.1.2.3 plugin and mBed algorithm. We chose alignments with pairwise identity (over 99.7%) and nucleotide length over 9,000 as consensus sequences. NCBI ORF finder with standard genetic code (https://www.ncbi.nlm.nih.gov/orffinder) was used to identify the coding sequence (CDS) and the polyprotein translation. *Iflavirus* consensus genomes are available in NCBI (BrBV_TN accession PV031990, IspV_TN PV023937, UeV_TN PV023938, UspV_TN PV023939).

### *Iflavirus* detection in SRA data

We searched available RNAseq datasets from aphid species in the Sequence Read Archive (SRA) available through NCBI (3 March 2024). We selected non-pea aphid experiments from field-collected aphids sequenced with the Illumina platform as paired-end reads and with mixed selection methods and chose one experiment per species (Table S4). The accession numbers (SRX) were used to obtain their corresponding run number (SRR) into a custom bash pipeline to download corresponding read files using the SRA Toolkit (https://github.com/ncbi/sra-tools/wiki). Read files submitted to NCBI as multiple interleaved files were downloaded as individual read files and concatenated into two separate files (R1 and R2) with a cat command line. We conducted CZ ID and TRAVIS analyses as above and used NCBI ORF finder to identify the CDS and the polyprotein translation. Host species taxonomy (>99.98% identity) from which we obtained viral sequences was confirmed by blasting reference COI barcode sequences (http://www.boldsystems.org/) against the SRA experiment set. The *Iflavirus* consensus genomes and polyprotein sequences obtained from SRA are available as Supplementary File 1

### *Iflavirus* screening via PCR

We used the primer design tool in Geneious Prime v.2024.0 to generate specific primer pairs targeting different regions encoding for the *Iflavirus* polyprotein of BrBV, IspV, UeV, and UspV (Table 2). Primer target specificities were confirmed by Sanger sequencing amplicons and aligning them to each *Iflavirus* genome. We also confirm the species specificity of each primer set by retesting all virus-positive parental samples (RNAseq data) against the entire virus panel (six primer sets). For *Iflavirus* screening, 800 μg of total RNA (extracted as above) was used for cDNA synthesis using iScript cDNA synthesis kit (Bio-Rad Laboratories, Inc., Hercules, CA, USA), which uses random hexamer primers. Then, 1 μL undiluted cDNA was used as a template in a 25 μL PCR reaction using Quick-Load Taq 2X Master Mix (New England Biolabs, Ipswich, MA, USA), 0.2 μM of each primer, and the following the thermal cycling conditions: 94°C for 5 min followed by 35 cycles of 94°C for 30 s, the corresponding annealing temperature for 30 s, 72°C for 30 s, and a final extension at 72°C for 3 min. We visualized PCR products in 1% agarose gels stained with SYBR Safe (Invitrogen, Inc).

**Table 2.**
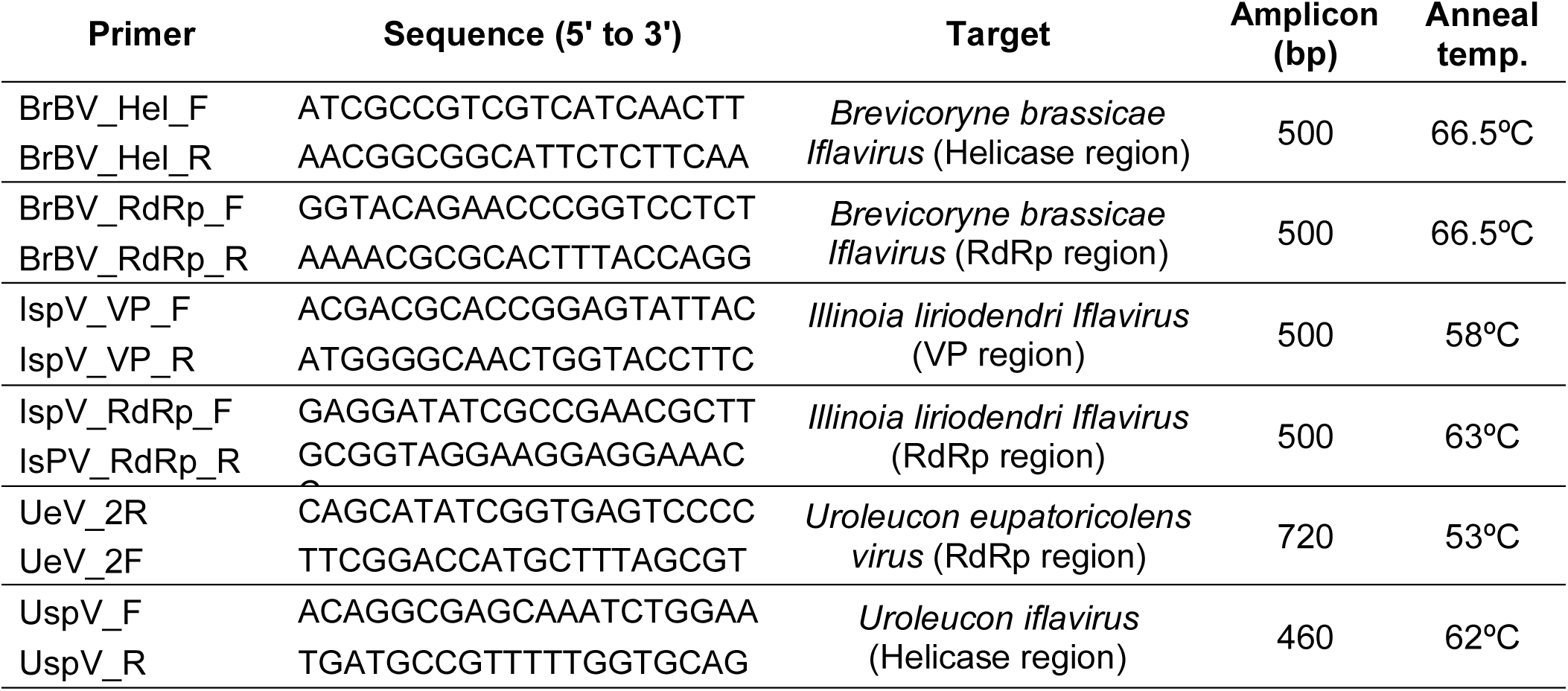
Primers used for aphid *Iflavirus* screening using conventional PCR.

### *Iflavirus* titer via qPCR

We used quantitative PCR (qPCR) to calculate relative virus titers with specific primers for BrBV and IspV (Table 3) and the aphid Glyceraldehyde 3-phosphate dehydrogenase (G3PDH) as an endogenous control gene for normalization of quantitative data and as an internal control of cDNA quality (73). We used 1 μL of undiluted cDNA (synthesized as above) as a template in a 20 μL reaction containing 1X PCR Buffer, Mg^2+^ (2 μM), dNTP mix (0.2 μM), and Taq (0.025 units/μl) (Invitrogen, Inc.) with 1X EvaGreen Dye (Biotium, Inc., Fremont, CA, USA) and the corresponding primer concentrations (Table 3). The qPCR parameters were an initial step of 95°C for 3 min and 40 cycles of 95°C for 10 s, 60°C for 30 s; followed by a melt curve (65°C followed by 95°C (5 s at 0.5°C/cycle). All samples were run in triplicates in 96-well plates (Bio-Rad, Inc.) with a Bio-Rad CFX96 System (Bio-Rad). Single peaks in melting curve analyses were used to confirm gene-specific amplification and rule out non-specific amplification and primer-dimer generation. Relative virus titers were calculated using the average Ct values and the comparative ΔCt (ΔCt = average endogenous control Ct values – average virus Ct values).

**Table 3.**
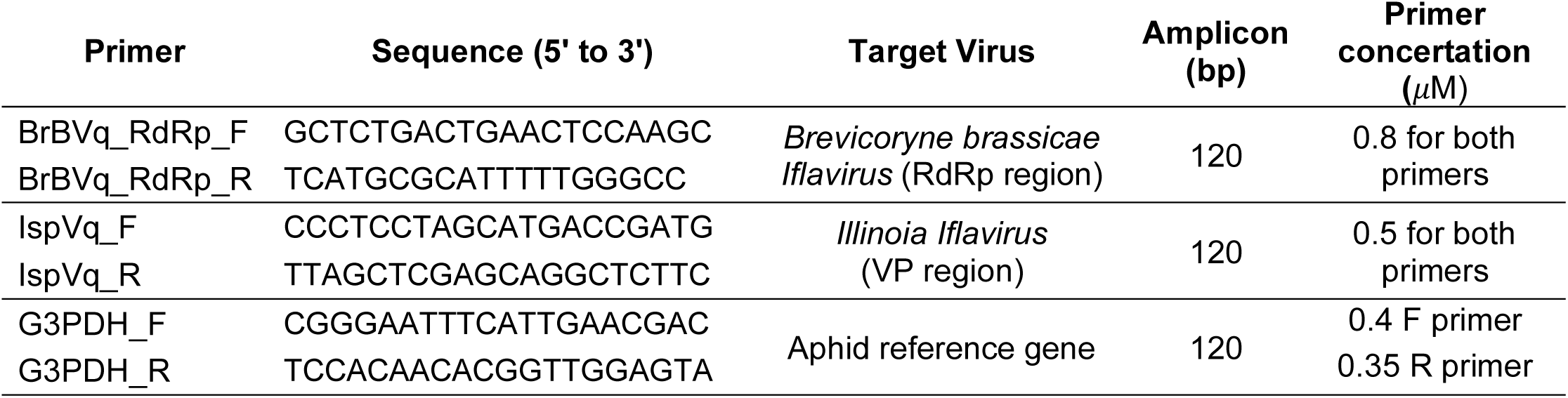
Primers were used to measure relative *Iflavirus* titers using qPCR.

### *Iflavirus* prevalence and heritability

To assess the prevalence of BrBV, IspV, UeV, and Usp in field populations, we screened single adult field-collected aphids (parental lines, P) using PCR (as above). We then assessed *Iflavirus* maintenance by PCR screening one to three individual adult aphids after one (F_1_) and three (F_3_) generations post-colonization using laboratory-grown plants.

For all subsequent transmission experiments, we used three lines of *Iflavirus*-positive aphids showing heritable infections for more than 30 generations in the laboratory [BrBV(+) *B. brassicae* aphids (lines A132 and A134) reared on cabbage (*Brassica oleracea*) and IspV(+) *I. liriodendri* aphids (line A76) reared on shepherd’s purse (*Capsella bursa-pastoris*)]. One *I. liriodendri* IspV(−) line (A65) reared on shepherd’s purse, and one *L. pseudobrassicae* BrBV(−) line reared on cabbage (A130) were used as negative controls. Unfortunately, no *Iflavirus*-negative *B. brassicae* lines were successfully colonized.

We assessed heritability as vertical transmission rates (VTR; percentage of infected females that transmit the virus to any of their offspring) and filial infection rates (FIR; percentage of infected offspring among all progeny from an infected mother) using two lines of BrBV(+) *B. brassicae* aphids (A132 and A134) and one line of IspV(+) *I. liriodendri* aphids (A76), totaling four biological replicates per line (reproductive females). We first placed individual 4^th^ instar nymphs in a Petri dish containing a detached leaf of the host plant and allowed them to reach parthenogenic reproduction. At 24 hours, the progeny of each female was immediately collected from the posterior ends of their mothers while being born and transferred to new detached leaves. The reproductive females that produced more than five nymphs were stored in Eppendorf tubes at −80°C and then screened for virus status by PCR, as above. The F_1_ nymphs of single reproductive females testing positive for *Iflavirus* infection were maintained on lab-grown plants for 10–14 days until reaching adulthood. Individual F_1_ adults were stored in Eppendorf tubes at −80°C until screened for virus status by PCR, as above. In addition, we used RNA obtained from eight individual *B. brassicae* adults (reproductive females from lines A132 and A134) to measure relative BrBV adult body titers (qPCR). Differences in relative titers (ΔCt) between BrBV(+) lines were analyzed using a two-tailed Mann-Whitney test (p < 0.05) using Data GraphPad Prism v.10.0 (www.graphpad.com).

### Within species transmission of iflaviruses

To investigate whether cohabitation enhances transmission rates of BrBV and IspV, we placed individual 4^th^ instar nymphs (*B. brassicae* and *I. liriodendri*, respectively) in a Petri dish containing a detached leaf of the appropriate host plant and allowed to reach parthenogenic reproduction. The reproductive females were allowed to cohabitate with their progeny for four days before removing all the F_1_ nymphs and placing them on fresh cabbage plants. Reproductive females and adult F_1_ aphids were individually placed in Eppendorf tubes at −80°C and later screened for virus status by PCR, as above. We conducted four biological replicates (reproductive females) per aphid line.

To determine whether Iflavirus was present in body secretions of persistently infected aphids to serve as a potential source of virus for horizontal transmission, we tested honeydew and cornicle secretions from *B. brassicae* adults for the presence of virus. We used forceps to apply gentle pressure to adults’ abdomens and collected honeydew directly from the aphid’s anus and cornicle droplets using glass capillary needles (volume 0.5 µL). Corresponding honeydew and cornicle droplets from several individual adults were pooled until obtaining 20–50 µL. Additionally, we tested plant leaf surfaces (contaminated with body secretions) where BrBV(+) aphid lines had been reared for 10 days. Individual cabbage leaves containing clusters of 8-12 aphids attached (nymphs and adults) were clipped, all aphids removed using forceps, and the leaves washed with 500 µL 1X phosphate-buffered saline (PBS) by pipetting up and down on the leaf surfaces. We extracted total RNA from body secretions and leaf-wash samples and tested by qPCR, as above. The cDNA of adult BrBV(+) aphids (whole-body, n=3) from which honeydew and cornicle droplets were collected served as a positive control for the qPCR.

To test *Iflavirus* oral ingestion results in active infection, we fed virus-free *I. liriondendri* aphids (line A65), on artificial aphid diets (74) supplemented with IspV(+) aphid homogenates. Whole-body aphid homogenates were prepared by high-speed shaking 10 adults in 1000 µL 1X PBS at 4°C with a Bead Mill Homogenizer (Omni, Kennesaw, GA, USA) for 4 min at 3.1 m/s followed by centrifugation at 12,000×g for 6 min at 4°C. The cleared supernatant was filtered with a 0.22 µm PDVF syringe filter and immediately used in feeding assays. We delivered the homogenate-supplemented insect diets (mixed 1:1) to 10–15 2^nd^ and 3^rd^ instar nymphs via parafilm sachets (400 µL). Homogenate-supplemented diets from the virus-positive aphid lines served as the infectious treatment, virus-negative lines served as negative controls, and a PBS-supplemented diet was used as the homogenate control. RNA was extracted from a 200 µL aliquot of the homogenate-supplemented diets, as above, to confirm the presence of IspV in the inoculum by PCR. To ensure only aphids that fed on the artificial diet were tested, we added 0.1% of blue food coloring (FD&C blue #1; Kroger Co., Cincinnati, OH, USA) to all diets, and only collected aphids whose abdomens were blue. Aphids were allowed to feed for 24 hours before being transferred to new plants for an additional seven days. We collected pools of 2–3 surviving aphids immediately after the feeding period (time 0), after two and seven days of being placed on host plants. All aphids from artificial diet assays were placed in Eppendorf tubes at −80°C. Four biological replicates of each feeding event were screened for virus by PCR, as above. *B. brassicae* aphids were excluded from all feeding assays due to the absence of colonized BrBV-free lines.

Since other aphid heritable viruses can systemically spread through the plant phloem (20, 41), we tested the potential for *Iflavirus* transmission by feeding on the host plant. we placed the roots of a small shepherd’s purse plant in an Eppendorf tube containing a 1:1 mixture of IspV(+) aphid homogenate and 1X PBS, secured the plant to the Eppendorf tube with parafilm, and placed the setup in a Petri dish. We then placed 2^nd^ and 3^rd^ instar nymphs from the IspV(−) *I. liriodendri* line on top of the leaves. Plants placed only in 1X PBS served as negative controls. To prevent surface contamination during the setup, plant leaves were washed with distilled water and a 5% bleach solution before placing the aphids. Aphids were allowed to feed on the leaves of plants whose roots were submerged in homogenates for three days, we then collected 1–3 surviving aphids and transferred the remaining surviving aphids to new unexposed plants for additional eight days to allow them to reach adulthood and parthenogenically reproduce on the virus-free plants. We collected all the surviving adult aphids (as individuals) and their progeny (in pools of 2–3 nymphs) produced on virus-free plants. Feeding dishes where all aphids died were excluded from the analysis. Aphids collected from six biological replicates (feeding setup) were placed in Eppendorf tubes at −80°C and until screened for IspV by PCR, as above.

### Cross-species transmission of iflaviruses

To determine whether iflaviruses are horizontally transmitted between aphid species from different genera known to cohabitate and feed on the same host plants, we kept three BrBV(+) *B. brassicae* and three BrBV(−) *L. pseudobrassicae* adults in a Petri dish containing a cabbage leaf disc for 48 hours. White waxy *B. brassicae* individuals were visually distinguished from green *L. pseudobrassicae*. Reproductive females of both species were collected into Eppendorf tubes at −80°C for subsequent testing for BrBV by PCR, as above. Progeny of the BrBV(−) *L. pseudobrassicae* adults were transferred to virus-free plants, allowed to mature, and tested for BrBV by PCR. Negative PCR results were re-tested by qPCR to confirm that negative results were not due to the PCR limit of detection. In total, we conducted four biological replicates (feeding setups).

To determine whether iflaviruses are horizontally transmitted cross-species by oral infection through the host plant, we placed a cabbage leaf steam in an Eppendorf tube containing 1:1 mixture of 1X PBS and BrBV(+) aphid homogenate (prepared as above). The leaf was secured to the Eppendorf tube with parafilm and washed with distilled water and 5% bleach before placing ten 2^nd^ and 3^rd^ instar nymphs of the BrBV(−) *L. pseudobrassicae* (line A130) on top of the leaves. Leaves placed in 1X PBS served as negative controls. Aphids were allowed to feed on the leaves submerged in homogenates for three days, then we collected 1–3 surviving aphids for testing by PCR and transferred the remaining survivors to new unexposed plants for additional eight days to allow them to reach adulthood and parthenogenically reproduce on the virus-free plants. Aphids collected from six biological replicates (feeding setups) were placed in Eppendorf tubes at −80°C until screened for BrBV by PCR and, as above, negative PCR results were confirmed via qPCR.

Last, to explore the BrBV potential to infect other species when all infection barriers are bypassed, we microinjected approximately 500 nL of undiluted BrBV(+) aphid homogenates in BrBV(−) 4^th^ instar *L. pseudobrassicae* nymphs. Aphid homogenates made with BrBV(−) *L. pseudobrassicae* adults and 1X PBS served as negative controls. After microinjection, aphids were reared on fresh cabbage plants. We collected ten live aphids one hour after the injection and at 3 and 7 days post-injection (dpi) in Eppendorf tubes, stored at −80°C, and screened for BrBV infections by qPCR. We also collected the F_1_ offspring produced by injected females as pools of 10 nymphs and tested them as above.

### *Iflavirus* titers in aphid tissues

To evaluate infection patterns of BrBV infection across various tissues, we briefly submerged adult *B. brassicae* aphids (10-14 days after emergence) in 70% ethanol and rinsed twice in 1X PBS before dissecting them under a stereomicroscope using sterile tweezers. Heads (including the salivary glands and brain), digestive tract (including bacteriocytes), developing embryos (including ovarioles), and the remaining carcass were separately collected. We pooled tissues of twenty individuals (maintained on the same plant) in 100 µL of PBS, seven biological replicates each, and stored at –80°C until processing and testing by qPCR, as above. Differences in relative titers between tissues (ΔCt) were analyzed using a Kruskal-Wallis test followed by multiple comparisons (p < 0.05) using Data GraphPad Prism v.10.0.

### Immunohistochemistry

We fixed adult BrBV(+) *B. brassicae* and BrBV(−) *L. pseudobrassicae* (negative controls) in 4% paraformaldehyde (Electron microscopy sciences, Hatfield, PA, USA) and stored at 4°C for approximately one week. Aphids were then placed in tissue cassettes (Thermo Fisher Scientific, Inc.), and processed through a standard dehydration/infiltration series before embedding in Ameraffin LP paraffin (Cardinal Health, Inc., Dublin, OH, USA) and stored at 4°C. We fixed aphid block sections (5 *μ*m) to charged microscope slides (Colorfrost Plus, Thermo Fisher Scientific, Inc.) by heating at 42°C overnight. Sections were put through a standard deparaffinization and rehydration series and incubated in 1X PBS for 10 min. Antigen retrieval was achieved by submerging sections in citrate-EDTA buffer (10 mM citric acid, 2 mM EDTA, 0.05% Tween20, at pH 6.2) at 88°C for 20 min and cooling down the sections at room temperature before blocking with 6% casein (Sigma-Aldrich, St. Louis, MO, USA). We then incubated the sections with a 1:200 dilution of a polyclonal rabbit anti-BrBV capsid peptide antibody (sequence RGDLEFKFVLNSNKFC) (GenScript, Piscataway, NJ, USA) at room temperature for 1 hour, followed by sequential incubation with biotinylated goat anti-rabbit, avidin-biotin alkaline phosphatase (VECTASTAIN ABC-AP Staining kit, Vector Laboratories, Inc., Newark, CA, USA), and Vector Red chromogen substrate (Vector Laboratories), according to manufacturer’s instruction. Between each incubation step, sections were rinsed with PBST (1X PBS, 0.1% Tween20) for 5 min. Tissues were counterstained with Meyer’s hematoxylin (Sigma-Aldrich), followed by 0.1% sodium bicarbonate (Sigma-Aldrich), and covered with CC/Mount (Sigma-Aldrich). Rothschild et al. (75), Forbes (76), and Simonet et al. (26) were used to confirm tissue types.

### *Iflavirus* influence on host fitness

To determine the effect of *Iflavirus* infection on life history traits in *I. liriodenri* lines [IspV(−) and IspV(+)], individual 4^th^ instar nymphs were isolated in a Petri dish containing a detached leaf of shepherd’s purse and allowed to reach parthenogenic reproduction. We assessed the effects of viral infection on fecundity as the number of offspring produced per female after four days across six biological replicates (reproductive females) per line. To determine the effects of viral infection on development times, we transferred the 1^st^ instar F_1_ nymphs (pools of 8–10) from single parents to new plants and recorded the number of days needed to reach parthenogenic reproduction (six biological replicates per line). We used a two-tailed Mann-Whitney test (p < 0.05) to analyze the differences in life history traits between lines using GraphPad Prism v.10.0. We did not test the effects of *Iflavirus* infection in *B. brassicae* due to the lack of BrBV(−) lines.

The production of winged or wingless offspring as a response to environmental conditions is an important factor in the spread of aphid-associated viruses (77). To measure the effects of viral infection on stress-induced wing induction, we randomly assigned 12-day-old *I. liriodenri* adults reared at low densities for three generations (< 10 aphids per plant) to Petri dishes (3.5 cm diameter) either as solitary treatment (one aphid per) or crowding treatment (12 aphids per dish) for 16 hrs. Aphids were then placed on a virus-free plant for three days and allowed to produce offspring. On day four, we removed the reproductive adults from the plant and allowed their offspring to reach adulthood. We recorded the percentage of offspring that developed across four biological replicates (treatment per line) and analyzed the differences using a two-tailed Mann-Whitney test (p < 0.05) in GraphPad Prism v.10.0. As before, we did not test wing induction responses in *B. brassicae* due to the lack of BrBV(−) lines.

Lastly, we evaluated if *Iflavirus* titers affect the titers of the obligatory bacterial symbiont, *Buchnera aphidicola*, due to co-localization in bacteriocytes. As above, we used 1 *μ*L of undiluted cDNA template in a 20 *μ*L qPCR reaction containing 0.5 *μ*M of GroES_F (5′ - CTTCGTCCGTTGCATGATCGT - 3′) and 0.8 *μ*M of GroES_R (5′ - TGCAGCAGAACCCGTAAAGAA - 3′), specific primers targeting the *B. brassicae Buchnera* gene GroEs (a cofactor essential for cell viability), and followed the same parameters established for *Iflavirus* titer. We used the cDNA of eight BrBV(+) adults that showed variable titers through previous experiments and simultaneously performed a qPCR for BrBV, *Buchnera,* and the aphid G3PDH (endogenous control). We calculated BrBV and *Buchnera* relative titers using the comparative ΔCt and analyzed the correlation between the relative titers of both symbionts with a Spearman Rank-Order test in Data GraphPad Prism v.10.0. Symbiont titer results for *I. liriodenri* lines are not included due to the lack of a species-specific *Buchnera* genome that would allow us to design high-efficiency qPCR primers.

### Phylogenetic analysis of iflaviruses

We aligned the polyprotein sequences of the newly and previously reported aphid iflaviruses (Table S6) and 37 reference *Iflavirus* species recognized by the ICTV (Table S9) in MAFFT v7.490 using FFT-NS-i_x1000 algorithm, matrix BLOSUM62, and 1.54 gap penalty (78). Then, used the polyprotein alignment to generate a Maximum–Likelihood tree using RaxML v8.2.12 with PROTGAMMAJTTF model and a majority rule bootstrap with 500 reps (79). To evaluate the consistency of clustering patterns among aphid iflaviruses, we also used polyprotein alignments (produced as above) to generate a Neighbor-Joining tree using the Jukes-Cantor distance model and 500 bootstrap replicates in Geneious Prime v2024.0 and an extended Maximum–likelihood tree that included 221 additional curated *Iflavirus* sequences (Table S10). Polyprotein sequences from *Dicistroviridae* (NP_066241.1 and NP_647481.1) and *Picornaviridae* (NP_041277.1 and NP_041007.1) served as outgroups in all analyses.

## Supporting information

Sequences

## Supplemental material

**Figure S1** Relative abundance of viral protein hits associated with different aphid species.

**Figure S2** Effect of *Iflavirus* infection on aphids’ fitness.

**Figure S3** Evolutionary analysis of aphid iflaviruses using a Neighbor-Joining tree.

**Figure S4** Extended maximum-likelihood analysis of aphid iflaviruses.

**Table S1** Metadata of field-collected parental aphids from which laboratory colonies were maintained in the laboratory for at least generations.

**Table S2** RNAseq metadata and CZ ID reads report.

**Table S3** Consensus genome characteristics of newly reported aphid *Iflavirus* sequences.

**Table S4** SRA datasets screened for *Iflavirus* reads.

**Table S5** *Iflavirus* polyprotein similarity matrix.

**Table S6** Sequences details of the newly and previously reported aphid iflaviruses.

**Table S7** Metadata of the field-collected aphids screened for *Iflavirus* infections.

**Table S8** *Iflaviridae* reference ORF database.

**Table S9** Reference *Iflavirus* species recognized by the ICTV.

**Table S10** Additional arthropod *Iflavirus* sequences used in the extended maximum-likelihood analysis.

**Supplementary File 1.** *Iflavirus* consensus genomes and polyprotein sequences obtained from SRA data sets.

## Data Availability

The aphid species barcode sequences are available through NCBI accession PV016919–PV01693. Sequencing raw reads are available through NCBI BioProject PRJNA1216975. *Iflavirus* genomes are available through NCBI accessions PV031990, PV023937, PV023938, PV023939.

## Acknowledgments

We thank Barbara S. Drolet at the Arthropod-Borne Animal Diseases Research Unit, USDA, ARS, Manhattan, KS, for providing insightful feedback on immunohistochemistry. Many thanks to Jessica Gutierrez at the USDA Arthropod-Borne Animal Diseases Research Unit and Taylor Do at the University of Tennessee Knoxville for technical assistance. We thank Dr. Heidi Goldrich-Blair, Keertana Tallapragada, and Meaghan Adler for providing insightful feedback on the project and manuscript. Special thanks to the East Tennessee farmers for logistical support during fieldwork. This work was funded by the National Science Foundation (NSF) award number DEB-2305653 to PRL and IOS-2152954 to BJP. BJP is a Pew Scholar in the Biomedical Sciences, funded by the Pew Charitable Trusts.

## Notes

### Competing Interest Statement

The authors have declared no competing interest.

