## Supplementary Figures and legends for "Heritable viral symbionts in the family *Iflaviridae* are widespread among aphids"

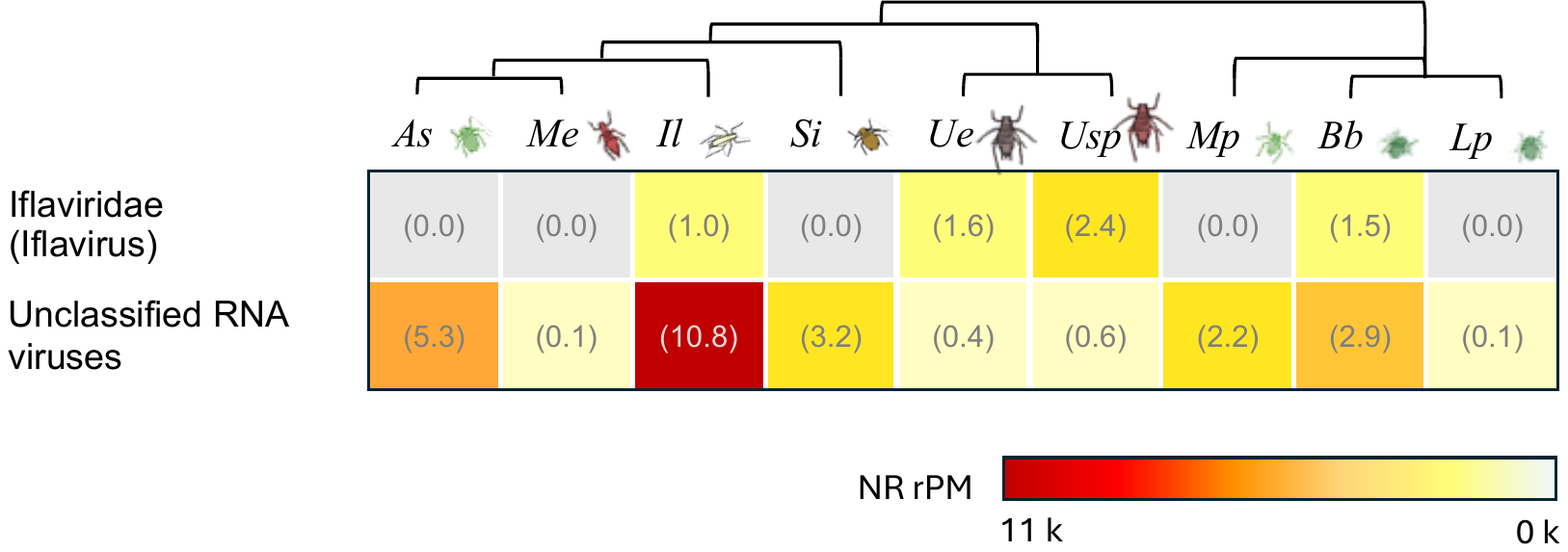

**Figure S1** Relative abundance of viral protein hits associated with different aphid species. NR rPM gradient bar indicates the proportion of reads aligning to iflaviruses and unclassified RNA viruses in the NCBI protein database (specific Nr rPM values in parenthesis). Aphid species depicted by their phylogenetic association (*Aulacorthum solani* (As), *Macrosiphum euphorbiae* (Me), *Illinoia liriodendri* (Il), *Sitobion ibarae* (Si), *Uroleucon eupatoricolens* (Ue), *Uroleucon* sp. (Usp), *Myzus persicae* (Mp), *Brevicoryne brassicae* (Bb), *Lipaphis pseudrobrassicae* (Lp).

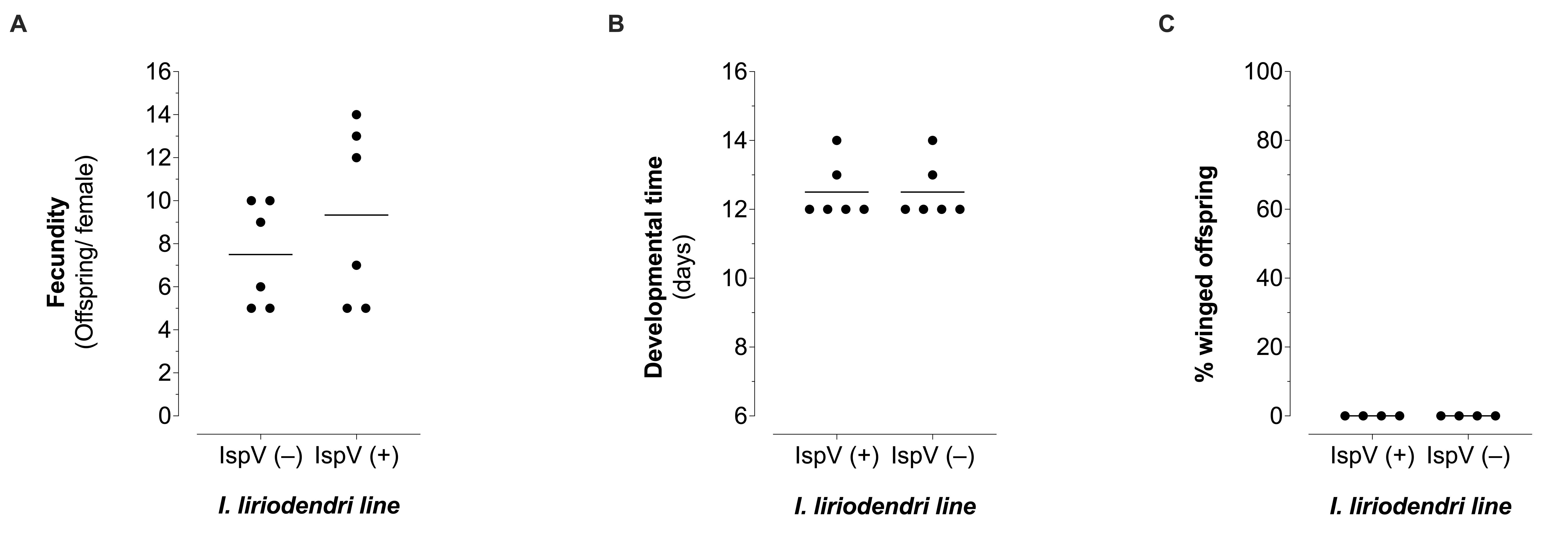

**Figure S2** Effect of *Iflavirus* infection on aphids’ fitness*.* The effects of viral infection on (**A**) Fecundity (number of offspring produced by an individual female in four days), (**B**) developmental time (number of days needed to reach parthenogenic reproduction after being born), and (**C**) stress-induced wing production (percentage of offspring that developed wings as adults when reproductive females were subject to solitary or crowding treatments) were evaluated in genetically independent *Illinoia liriodenri* lines [IspV(−) and IspV(+)].

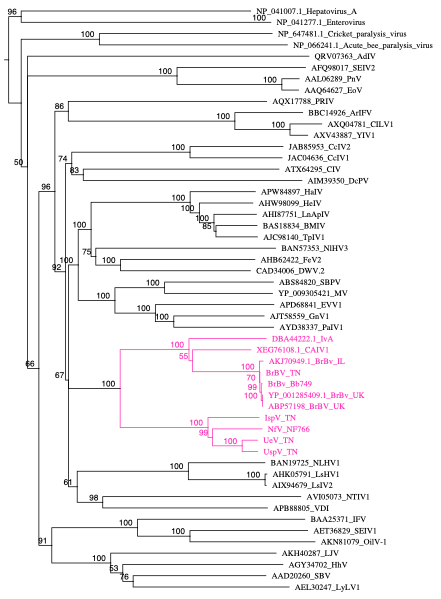

**Figure S3** Evolutionary analysis of aphid iflaviruses using a Neighbor-Joining tree. The tree was constructed with the Jukes-Cantor genetic distance model based on an MAFFT alignment of the polyprotein sequences of aphid Iflaviruses (pink clade) and ICTV-recognized members of Iflaviridae. Polyprotein sequences from *Dicistroviridae* (*Acute bee paralysis virus*, NP_066241.1 and *Cricket paralysis virus*, NP_647481.1) and *Picornaviridae* (*Enterovirus* C, NP_041277.1, and *Hepatovirus* A, NP_041007.1) served as outgroups. Bootstrapping values on each node represent the percent from 500 replicates.

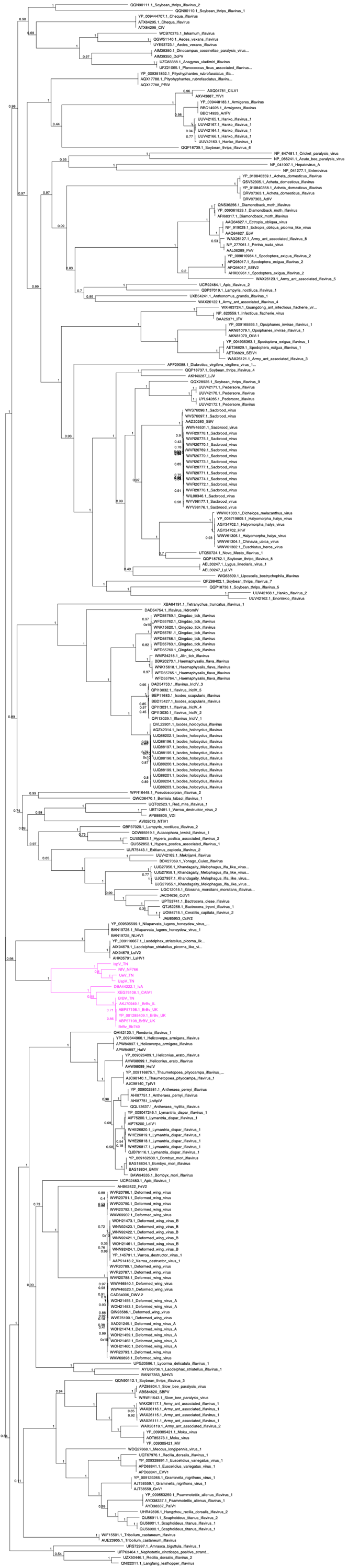

**Figure S4** Extended maximum-likelihood analysis of aphid iflaviruses. The tree was constructed using the Maximum-likelihood method with the protein gamma JTT matrix based on an MAFFT alignment of the polyprotein sequences of aphid iflaviruses (pink clade) and sequences from a wide range of iflaviruses infecting arthropods. Polyprotein sequences from *Dicistroviridae* (*Acute bee paralysis virus*, NP_066241.1 and *Cricket paralysis virus*, NP_647481.1) and *Picornaviridae* (*Enterovirus* C, NP_041277.1, and *Hepatovirus* A, NP_041007.1) served as outgroups. Bootstrapping values on each node represent the proportion (0–1) from 500 replicates.
